# Differential Effects of Distinct Type I Interferons and IFN Inducers on Fc Receptor landscape and myeloid cell function

**DOI:** 10.64898/2026.10.01.756027

**Authors:** Rémi Bru, Océane Paris, Guillaume Bossis, Stephanie Boireau, Mar Naranjo-Gomez, Mireia Pelegrin

## Abstract

Antiviral monoclonal antibodies (mAbs) rely on Fc-mediated effector functions, which depend on interactions with Fc receptors (FcRs) expressed on immune cells such as monocytes and dendritic cells (DCs). While combinations of antiviral mAbs with type I interferons (IFN-I) or IFN inducers are being explored to enhance therapeutic efficacy, the impact of these immunomodulators on FcR expression remains poorly understood. In this study, we investigated the effects of IFN-α2, IFN-β, and the IFN-inducer TAK-981 (a SUMOylation inhibitor) on FcR modulation and functional activation in human monocyte subsets and monocyte-derived DCs (moDCs). Using flow cytometry and imaging techniques, we mapped the expression of activating FcγRs (CD64, CD32A, CD16A) and neonatal FcR (FcRn) under steady-state and stimulated conditions. We found that IFN-I selectively downregulates CD16A expression in classical monocytes while preserving other FcRs, whereas TAK-981 reduces FcRn expression in classical monocytes without affecting CD16A. Additionally, IFN-β and TAK-981 but not IFN-α2, selectively enhanced FcγRI expression in moDC. IFN-β also induced a stronger maturation and functional activation program in monocytes and moDCs compared with IFN-α2, whereas TAK-981 largely reproduced the IFN-β-driven phenotype and secretory profile. Furthermore, we demonstrated that IFN-I signaling alters the spatial distribution of FcRs, with IFN-β and TAK-981 promoting distinct redistribution patterns of FcγRs in monocytes and moDCs. These findings reveal a novel layer of FcR regulation, highlighting that IFN-I and IFN inducers differentially modulate FcR expression, localization, and functional activation in a cell-type- and stimulus-specific manner. Overall, this work provides a framework for understanding how IFN-I-based immunotherapies may influence Fc-mediated immune functions while preserving FcRn-dependent antibody persistence.

## INTRODUCTION

Antiviral monoclonal antibodies (mAbs), including broadly neutralizing antibodies (bNAbs), have emerged as a promising therapeutic strategy against severe viral infections ^1–10^. Several mAbs are already approved for clinical use and many others are currently being tested in clinical trials, notably for HIV-1 and HBV infection ^2,4,7,11^. Their therapeutic effect relies not only on the neutralizing activity of their Fab domain through binding to viral antigens, but also on Fc-mediated effector functions, which involve interactions with Fc receptors (FcRs) expressed on immune cells (e.g., natural killer cells, monocytes, and dendritic cells (DC)) as well as activation of the complement system ^12,13^. Fc–FcR interactions are highly complex, involving multiple activating and inhibitory Fc gamma receptors (FcγRs) with distinct affinities for different IgG subclasses, the antibody isotype most commonly used in clinical practice. In addition, the neonatal Fc receptor (FcRn), which also binds IgG, plays a central role in regulating antibody biodistribution and serum half-life. FcRs are differentially expressed across immune cell populations, enabling a broad range of effector functions ^12^. Beyond direct viral neutralization, antiviral mAbs mediate Fc-dependent effector functions, including antibody-dependent cellular phagocytosis (ADCP), antibody-dependent cellular cytotoxicity (ADCC), and complement-dependent cytotoxicity (CDC). By forming immune complexes with viral antigens, they also engage FcγRs on innate immune cells, promoting antigen uptake and presentation, cytokine production, and the induction of antigen-specific protective immunity (vaccinal effects) ^14–17^. The therapeutic potential of antiviral mAbs can be enhanced through Fc engineering, which modulates antibody pharmacokinetics and Fc-mediated immune functions. In particular, mutations that increase binding to activating FcγRs, (i.e. GAALIE, GASDIE, and GASDALIE among others)^18–20^ potentiate Fc-enhance Fc-dependent effector functions (reviewed in ^21^), whereas FcRn-binding mutations, such as the LS mutation, prolong serum half-life^22^. These engineering strategies have been shown to improve the *in vivo* efficacy of antiviral mAbs ^20,23^.

Given the pivotal role of Fc-FcγR interactions in mediating antiviral immunity, modulation of FcR expression—whether by immunomodulatory molecules or inflammatory conditions associated with viral infections— can significantly influence the therapeutic efficacy of antiviral mAbs. We previously demonstrated in a mouse model of retroviral infection that pro-inflammatory cytokines and the inflammatory environment modulate FcR expression in myeloid cells, including DC, monocytes and neutrophils which are key mediators of protective immunity induced by antiviral mAbs ^24–26^. We further showed that the inflammatory environment associated with HIV-1 infection modulates FcR expression in myeloid cells ^27^. Despite these findings, modulation of FcR expression remains largely overlooked in most preclinical and clinical studies evaluating antiviral mAbs, either as monotherapies or in combination strategies.

Several preclinical studies and ongoing clinical trials are currently investigating the potential synergistic benefits of combining antiviral mAbs with type I interferons (IFN-I) or IFN-inducers to enhance therapeutic efficacy ^28–30^. However, the effect of IFN-I/IFN-I inducers on FcR modulation are still poorly understood. This is a critical consideration, as modulation of FcR expression in the context of combination therapy could significantly impact the therapeutic efficacy of antiviral mAbs. While combinations of IFN-I/IFN inducers with mAbs have shown improved efficacy in certain viral infections, this effect is not systematically observed and appears to depend on the specific virus, preclinical model, and mAb tested ^28–30^. In the particular case of HIV-1 infection, bNAbs are currently being tested in two clinical trials combined with either IFN-α2 (NCT03588715) or the TLR9 agonist (lefitolimod)^28^. However, the clinical benefit of these combinations has yet to be clearly demonstrated. While there are multiple subtypes of IFN-I, including 13 IFN-α subtypes, IFN-β, IFN-ω, IFN-ε, and IFN-κ ^31^, IFN-α2 remains the most widely used in clinical settings. Nevertheless, preclinical and experimental evidence suggests that specific IFN-α subtypes and IFN-β may more effectively enhance antiviral immune responses and control viral propagation ^32–36^, suggesting that the choice of IFN-I subtype in combinatorial therapeutic approaches mighty be a key factor in achieving optimal protection. Other IFN-I inducers, such as STING agonists (i.e.2’3’-cGAMP, DiABZI)^37–39^, have also gained increasing interest for their potential to enhance antiviral immune responses and have been explored in combination therapies with antiviral mAbs. Among novel IFN-I inducers, the SUMOylation inhibitor TAK-981 (Subasumstat) has also emerged as a promising candidate for combinatorial approaches with mAbs. Indeed, TAK-981 promotes the production of IFN-β and has been tested in Phase I clinical trials for cancer (NCT03648372, NCT04074330). TAK-981, has been shown to reactivate IFN-I signaling, leading to upregulation of IFN-I gene expression, DC activation, and enhanced NK cell and macrophage functions^40–42^. Preclinical studies in mice demonstrate that TAK-981 enhances the efficacy of therapeutic antibodies (e.g., rituximab, daratumumab) suggesting potential applications in antibody-based immunotherapies ^43,44^. However, its role in the modulation of FcR in human myeloid cells has not been thus far studied and the knowledge in the direct cell-intrinsic effects of TAK-981 on specific human myeloid cells (i.e. monocytes and DC) in terms of functional activation is still limited ^41,45^.

In this study, we aimed to investigate the effects of IFN-I subtypes (IFN-α2, IFN-β) and the IFN-I inducer TAK-981 on the functional activation of human monocyte subsets and monocyte-derived DCs (moDCs), with a particular focus on the modulation of FcR expression. DCs are a diverse group of antigen-presenting cells vital for starting and directing immune responses. While all DCs can take up, process, and present antigens to T cells, different DC types vary in their origins, locations, migration, and specific immune functions. MoDCs, often called “inflammatory moDCs”, are particularly important during inflammation and infection ^46^ ^47^. These cells are thought to cooperate with conventional DCs, playing a complementary role that amplifies responses to inflammatory or infectious stimuli. In terms of FcR repertoire, MoDCs show a strong expression of FcγRIIA (CD32A), low levels of FcγRIIIA (CD16A), very low or absent expression of FcγRI (CD64) and high levels of the FcRn. Beyond DCs, other myeloid cells, notably monocytes, are also critical in the orientation of immune responses when activated by ICs through their interaction with FcRs ^15,26^. Monocytes can be segregated by CD14 and FcγRIII (CD16) expression levels into 3 distinct subsets. Non classical monocytes express high levels of CD16 (CD14^+^CD16^++^), intermediate monocytes express high levels of CD14 and CD16 (CD14^++^CD16^++^) and are functionally distinguishable from classic CD14^+++^CD16^+^ monocytes based on homing marker expression, antigen presentation capabilities and levels of proinflammatory cytokines produced after activation by different stimuli ^48^. Monocytes are recruited to sites of infection, where they actively contribute to the inflammatory process. The different monocyte subsets express a distinct repertoire of FcRs, including the activating receptors FcγRI (CD64), FcγRIIA (CD32A) and FcγRIIIA (CD16A) as well as FcRn^12^.

Given the clinical use of Fc-engineered bNAbs with enhanced affinity for different FcRs, in the herein study we mapped the expression landscape of activating FcγRs (CD64, CD32A, CD16A) and FcRn in monocyte subsets and moDC under steady-state conditions and upon IFN-I/IFN inducer activation. Our findings provide new insights into the differential effects of IFN-α, IFN-β and the IFN-I inducer TAK-981 on FcR modulation in monocyte subsets and moDCs. Additionally, we present a comprehensive analysis of the direct effect of TAK-981 on the functional activation of monocytes and moDCs in terms of phenotypic activation and secretion profiles. Overall, our research provides new insights into how specific IFN-I/IFN-I inducers differentially modulate the expression, cellular localization, and distribution patterns across monocyte subsets and moDcs, while also shaping their cytokine secretion profiles in a molecule-specific manner. Notably, IFN-I signaling was associated with distinct changes in FcR distribution patterns, with more pronounced spatial redistribution observed in moDCs. Together, these findings highlight that IFN-I signaling and pharmacological IFN-I induction regulate not only the abundance of FcRs but also their cellular positioning and distribution patterns, revealing an additional layer of FcR regulation that differs according to both cell type and stimulus. These insights should be taken into account to enhance the efficacy of mAb-based antiviral therapies, including Fc-engineered antibodies, when used in combination with IFN-I or IFN-I inducers.

## RESULTS

### IFN-I selectively reduce CD16A expression while preserving FcRn in human monocytes

First, we mapped the expression landscape of activating FcγRs (CD64, CD32A, CD16) and FcRn on human monocyte subsets under steady state conditions. Monocytes were isolated from blood samples from healthy donors (HD) using a magnetic bead-based negative selection, allowing purification of untouched monocytes (**Figure 1A**). This approach enables the isolation of the three monocyte subsets, which are identified based on their differential expression of CD14 and CD16 (**Figure 1B**). Among these, classical and non-classical (NC) monocytes represented the predominant populations (**Figure 1C**) and were therefore selected as the focus of subsequent analyses. Consistent with their phenotypic identity, NC monocytes expressed higher levels of CD16A. At steady state condition, classical monocytes exhibited higher expression of the high-affinity FcγR CD64, whereas NC monocytes showed elevated FcRn expression, as already described ^49^. Both monocyte subsets, however, exhibited comparably high expression of CD32A (**Figure 1D**). We next investigated whether IFN-I modulate FcR expression using two clinically relevant IFN-I subtypes, IFN-α2 and IFN-β. IFN-α2 is widely used therapeutically, whereas IFN-β has been reported to induce more potent antiviral responses and more effectively restrict viral dissemination^32–36^. Optimal stimulation conditions were established by dose-response analyses, selecting concentrations that induced maximal expression of the IFN-responsive marker CD317 while eliciting comparable signaling responses for both IFN-I subtypes (**Supplementary Figure 1**). Neither IFN-α2 nor IFN-β significatively modulated the expression of the activating FcγRs CD32A and CD64 or the FcRn on monocyte subtypes. On the contrary, both types of IFN-I lead to reduced expression of CD16A on classical monocytes (**Figure 1D**). Together, these findings demonstrate that IFN-I selectively downregulate CD16A while preserving the expression of other activating FcγRs and FcRn, revealing a targeted remodeling of the FcR repertoire in human monocytes.

**Figure 1.**
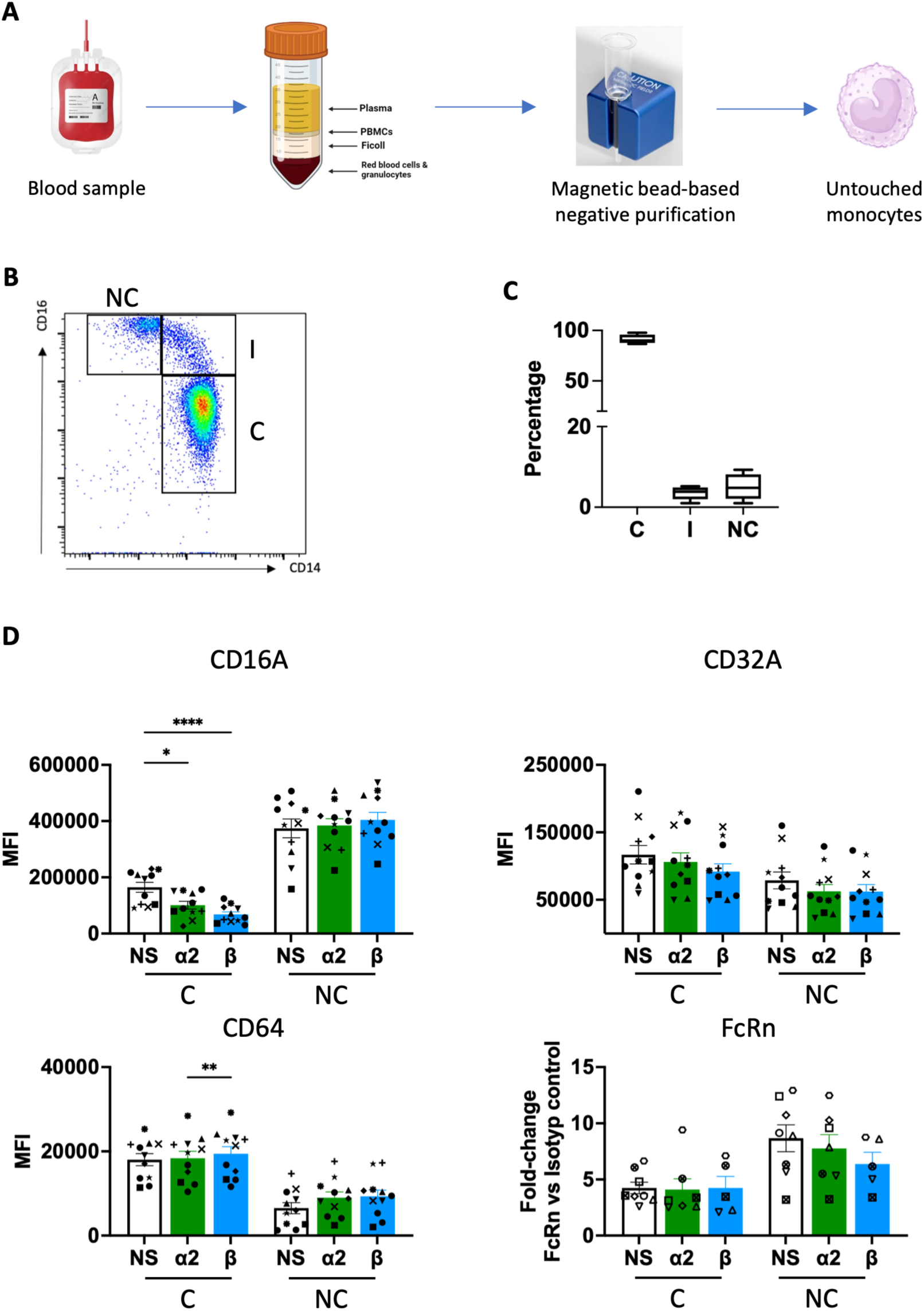
FcR expression on monocyte subsets. (A) Monocytes from healthy donors were purified from blood samples by magnetic bead-based negative selection. (B). Gating strategy used for identification of monocyte subsets. (C) Frequencies of monocyte subsets. (D) FcR expression on monocyte subsets upon activation by IFN-I, assessed by spectral flow cytometry. For FcRn, data are MFI of anti-FcRn monoclonal antibody and isotype MFI ratio. Values are mean +/- SD of at least 11 for FcγR and 5 for FcRn (D), independent measurements (different symbols represent data from different HD samples). Statistical analysis was performed by Normality and RM one-way ANOVA or Friedman test. C: classical monocytes; NC: non-classical monocytes, NS: non-stimulated. α2: stimulation with IFN-α2 at 10 000 U/mL; β: stimulation with IFN-β at 1 nM. Significance was assigned as follows: ∗p <0,05 ∗∗p <0,01 and ∗∗∗p < 0.001.

#### Comparable phenotypic and functional activation of classical and NC monocytes by IFN-α2 and IFN-β, though IFN-β induces specific quantitative differences in their activation profiles

We next assessed the phenotypic and functional activation of classical and NC monocytes following stimulation with IFN-α2 and IFN-β. To characterize phenotypic changes, we assessed the expression of cell surface markers associated with T-cell co-stimulation and antigen presentation (CD86, HLA-DR), cell adhesion (CD62L), and chemotaxis (CXCR4 and CCR5) (**Figure 2A**). Both IFN-I subtypes significantly increased CD86 expression on NC monocytes, whereas a significant increase in classical monocytes was observed only following IFN-β stimulation. Similarly, both IFN-I subtypes upregulated CD62L expression in classical and NC monocytes, although the increase induced by IFN-β was significantly greater than that induced by IFN-α2 in NC monocytes. In contrast, IFN-I stimulation led to decreased levels of the chemokine receptors CXCR4 and CCR5. Specifically, IFN-β induced a more pronounced downregulation of CXCR4, significantly reducing its expression in both classical and NC monocytes compared with either unstimulated or IFN-α2-treated cells. Likewise, IFN-β selectively decreased CCR5 expression in classical monocytes, whereas IFN-α2 had no significant effect.

**Figure 2.**
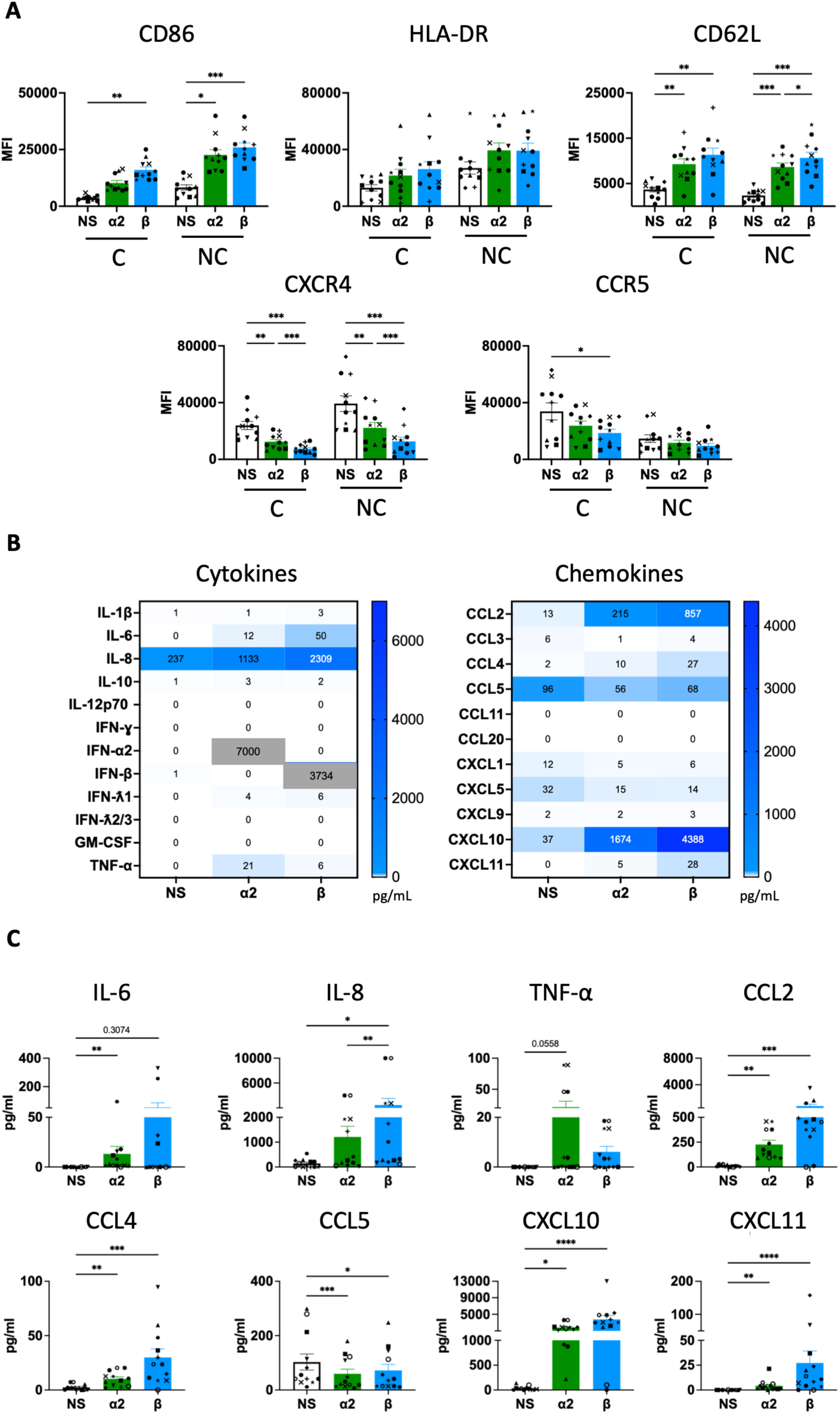
Activation of monocyte subsets by IFN-I subtypes. Monocytes from healthy donors were purified from blood samples by magnetic bead-based negative selection and activated with IFN subtypes for 18h. (A) Phenotypic activation of monocyte subsets assessed by monitoring cell surface markers, by spectral flow cytometry. (B-C) Cytokine and chemokine secretion profile of whole monocyte populations, represented as a heatmap (B) and histograms (C). The secretion of cytokines and chemokines was measured in pg/mL by LegendPlex assay. Values are mean +/- SD of at least 11 (A) and 12 (B-C), independent measurements (different symbols represent data from different HD samples). Statistical analysis was performed by Normality and RM one-way ANOVA or Friedman test. C: classical monocytes; NC: non-classical monocytes, NS: non-stimulated. α2: stimulation with IFN-α2 at 10 000 U/mL; β: stimulation with IFN-β at 1 nM. Significance was assigned as follows: ∗p <0,05 ∗∗p <0,01 and ∗∗∗p < 0.001.

To determine whether these phenotypic changes were accompanied by functional activation, we next analyzed the cytokine and chemokine secretion profile of IFN-I-stimulated monocytes (**Figure 2B**). Both types of IFN-I led to increased secretion of IL-8, CCL2, CCL4, CXCL10 and CXCL11. Overall, the secretory profiles induced by the two IFN-I subtypes were highly similar, with IL-8 representing the only mediator produced at significantly higher levels following IFN-β stimulation as compared to IFN-α2. IL-6 was also observed in a fraction of IFN-α2 and IFN-β-activated monocytes, although significant differences were only observed in IFN-α2-activated cells due to the high sample variability observed upon IFN-β activation. A similarly high degree of variability was observed for TNF-α secretion. Overall, these results provide a comparative and comprehensive study of phenotypic and functional activation of monocytes by IFN-I subtypes. They demonstrate that IFN-α2 and IFN-β elicit broadly comparable phenotypic and functional activation programs in human monocytes. Nevertheless, IFN-β consistently induced stronger quantitative changes in selected phenotypic markers and inflammatory mediators, indicating that it amplifies, rather than qualitatively alters, the IFN-I response.

### The IFN-I inducer TAK-981 partially recapitulates IFN-β-mediated modulation of monocyte phenotype and function

We next investigated the effect of TAK-981 on monocyte activation. This SUMOylation inhibitor has been reported to induce IFN-β production and exert immunomodulatory functions ^40–42^. However, its direct effects on the phenotypic and functional activation of distinct monocyte subsets remain poorly characterized ^41,45^. We first assessed the effect of this IFN-I inducer on FcR expression. Dose-response analyses allowed us to select the optimal TAK-981 concentration that induced maximal expression of the IFN-responsive marker CD317 and elicited comparable signaling responses than those observed upon IFN-α2 and IFN-β activation (**Supplementary Figure 1**). TAK-981 activation of monocytes did not affect the expression of CD32a and CD64 (**Figure 3A**), similarly to IFN-β (**Figure 1E**). However, unlike IFN-β, TAK-981 did not decrease the expression of CD16A in either monocyte subsets. Instead, it selectively decreased in FcRn expression in classical monocytes (**Figure 3A**), indicating that IFN-I induction by TAK-981 does not fully reproduce the FcR remodeling elicited by recombinant IFN-β. We next evaluated the effects of TAK-981 on monocyte phenotype. Treatment with TAK-981 increased the expression of CD86 and CD62L in both classical and NC monocytes, enhanced HLA-DR expression selectively in NC monocytes, and reduced CXCR4 expression in both subsets as well as CCR5 expression in classical monocytes (**Figure 3B**). These changes closely resembled those induced by IFN-β. Finally, we assessed the secretory response of TAK-981-treated monocytes (**Figure 3C**). TAK-981 promoted the secretion of IL-6, TNF-α, IL-8, CCL2, CCL4, CXCL10, and CXCL11. Overall, the cytokine and chemokine profile largely mirrored that induced by IFN-β. Collectively, these findings demonstrate that TAK-981 largely recapitulates the phenotypic and functional activation program induced by IFN-β while exhibiting a distinct pattern of FcR regulation, particularly with respect to CD16A and FcRn expression.

**Figure 3.**
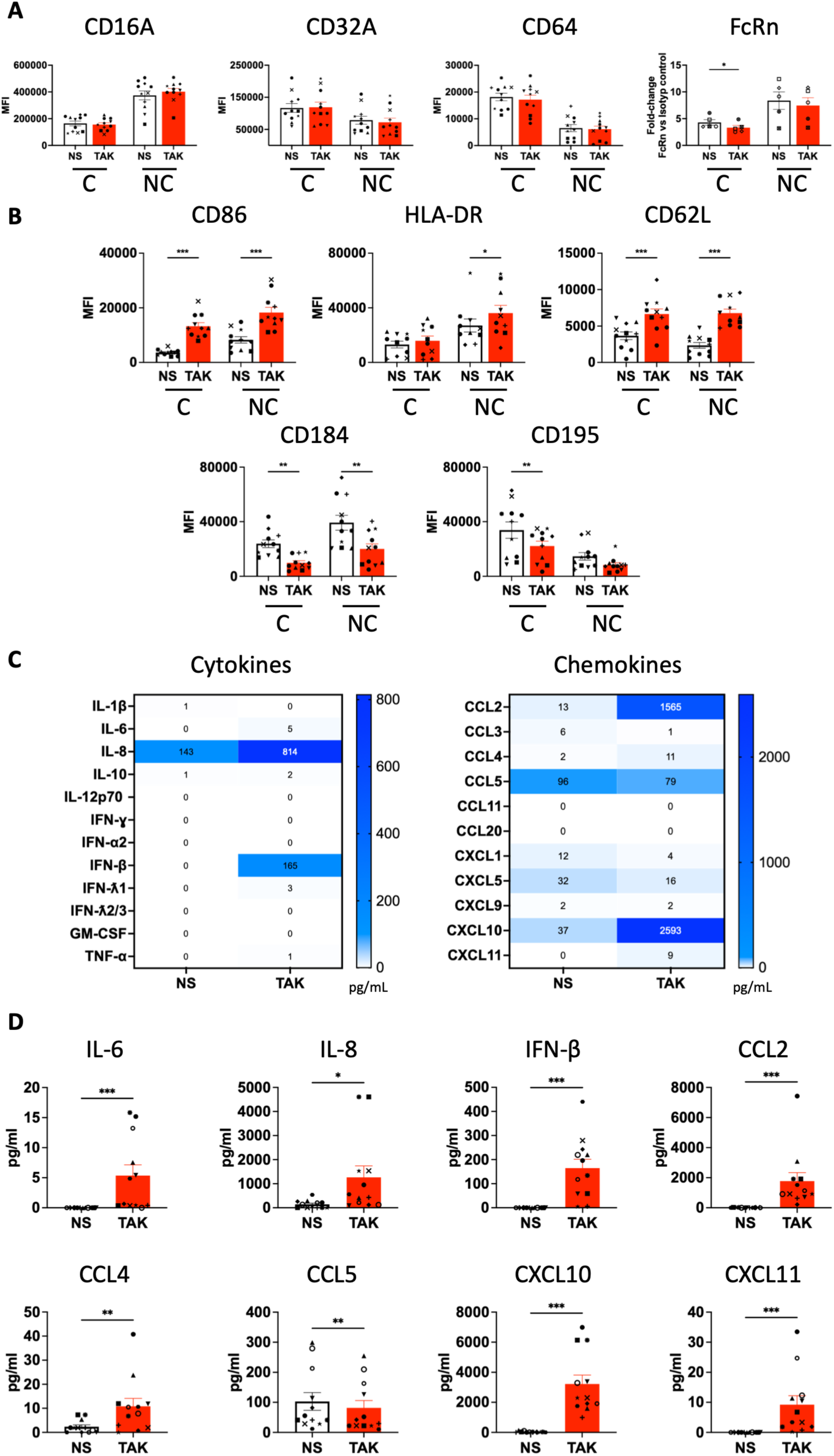
Activation of monocyte subsets by TAK-981. Monocytes from healthy donors were purified from blood samples by magnetic bead-based negative selection and activated with TAK-981 at 1 µM for 18h. (A) FcR expression on monocyte subsets, assessed by spectral flow cytometry. For FcRn, data are MFI of anti-FcRn monoclonal antibody and isotype MFI ratio. (B) Phenotypic activation of monocyte subsets assessed by monitoring cell surface markers, assessed by spectral flow cytometry. (C-D) Cytokine and chemokine secretion profile of whole monocyte populations, represented as a heatmap (C) and histograms (D). The secretion of cytokines and chemokines was measured in pg/mL by LegendPlex assay. Values are mean +/- SD of 11 (A), 10 (B) and 12 (C-D), independent measurements (different symbols represent data from different healthy donor (HD) samples). Statistics were performed by Normality and paired t test or Wilcoxon test. C: classical monocytes; NC: non-classical monocytes, NS: non-stimulated. independent measurements. Significance was assigned as follows: ∗p <0,05 ∗∗p <0,01 and ∗∗∗p < 0.001. Significance was assigned as follows: ∗p <0,05 ∗∗p <0,01 and ∗∗∗p < 0.001.

### IFN-I signaling modifies the spatial distribution patterns of FcR in human monocytes

We next examined whether IFN-I signaling alters the spatial organization of FcRs receptors in human monocytes. Using ImageStream®, which combines flow cytometry with high-resolution cellular imaging, we assessed FcR expression and localization at the single-cell level. ImageStream analysis provided additional insights on IFN-I- and TAK-981-induced modulation of FcR dynamics, notably on their cellular distribution (**Supplementary figure 2**). In particular, CD16A displayed a predominantly polarized distribution toward one pole of the cell under basal conditions, whereas IFN-α2 and IFN-β induced a more dispersed, punctate distribution of CD16A throughout the cell. To further characterize the spatial organization of FcRs, we used the Similarity Index -which quantifies the similarity of FcR fluorescence intensity distributions-to compare their fluorescence intensity patterns. Under basal conditions, most FcR pairs exhibited low levels of similarity, indicating distinct spatial patterns within the analyzed cellular compartment (**Figure 4A).** In contrast, FcRn and CD32A yielded a higher Similarity Index, suggesting that the two receptors exhibit similar spatial distributions and may be in close spatial proximity.

**Figure 4.**
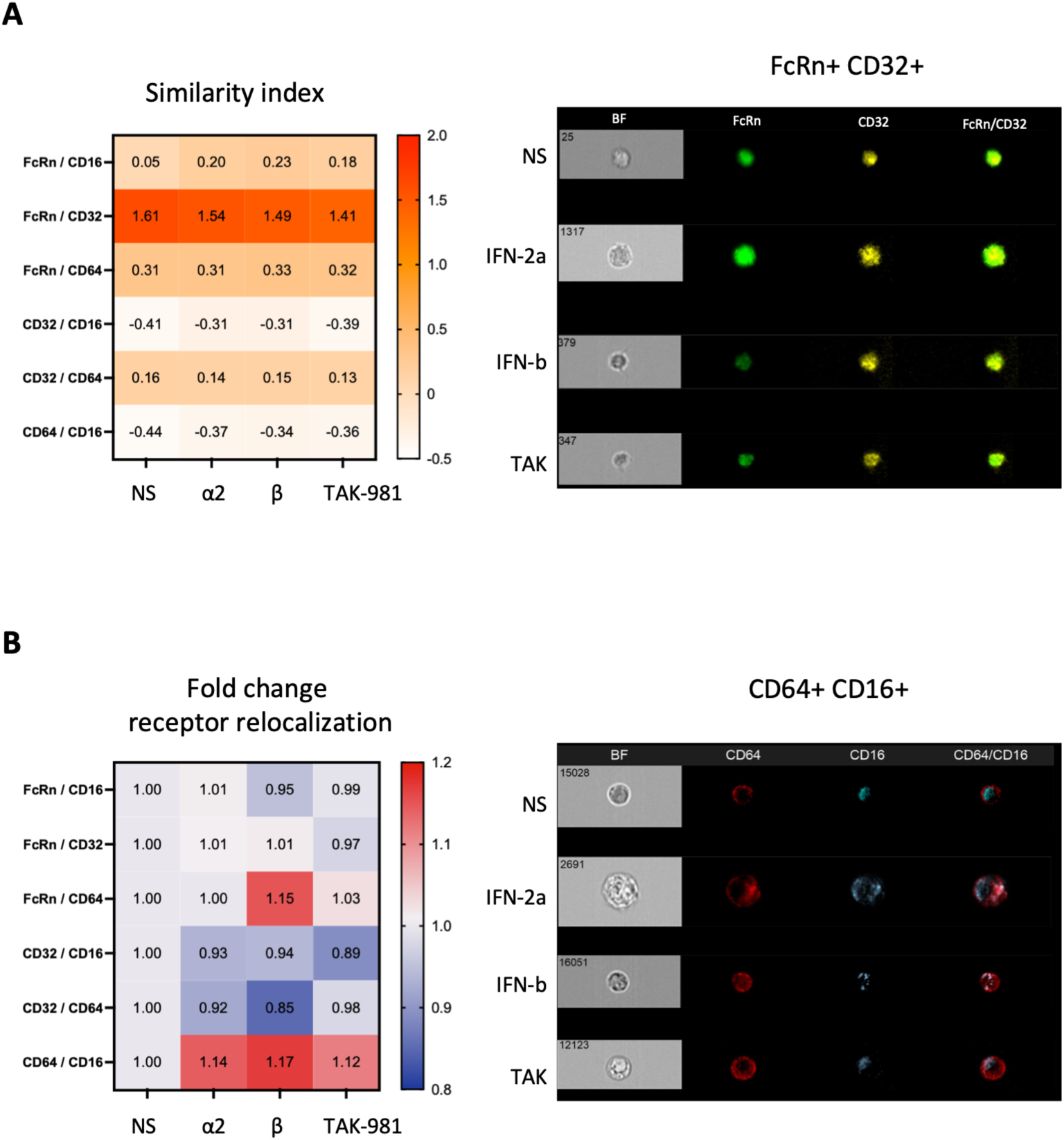
Cellular localization of FcR upon activation of monocyte by IFN-α2, IFN-β and TAK-981. Monocytes from healthy donors were purified from blood samples by magnetic bead-based negative selection and activated by IFN-α2 at 10 000 U/mL, IFN-β at 1 nM or TAK-981 at 1 µM for18h. (A) Similarity index heatmap for the different receptors obtained via manual compartmentalization using IDEAS® software. Fluorescence images show respective receptor expression and their merge after stimulation for the receptor pair with the highest level of co-expression (n = 3). (B) Fold change heatmap of Bright Detail Similarity R3 (BDS) scores calculated using the IDEAS® automated wizard to quantify colocalised receptor expression. Images are representative of three independent donors.

To determine whether IFN-I signaling altered FcR distribution patterns, we quantified changes in the receptor relocalization index following stimulation. This analysis revealed stimulus-dependent changes in the distribution of specific receptor pairs. IFN-β was associated with increased relocalization of FcRn and CD64, whereas IFN-α2, IFN-β, and TAK-981 promoted changes in the distribution of CD64 and CD16A (**Figure 4B**). Importantly, these changes did not reflect a uniform redistribution of FcRs. While IFN-α2 and IFN-β induced a more pronounced transition of CD16A from its polarized pattern toward a more dispersed, punctate distribution throughout the cell, TAK-981 produced a pattern more similar to the basal organization, with CD16A remaining predominantly polarized. Thus, although IFN-I stimulation and TAK-981 affected FcR distribution, they appeared to promote distinct patterns of receptor organization.

Together, these findings suggest that IFN-I signaling not only modulates FcR expression but may also remodel their spatial organization at cellular level, potentially influencing FcR cooperation and downstream signaling.

### IFN-I subtypes differentially remodel FcγR expression in moDC, with IFN-β promoting CD64 upregulation and CD32A downregulation

To investigate whether IFN-I signaling influences FcR in expression in moDC, CD14+ monocytes were purified from HD blood samples by magnetic bead-based positive selection and differentiated into moDCs by culturing them for 6 days in the presence of GM-CSF and IL-4 (**Figure 5A**). We first characterized the expression profile of activating FcγRs and FcRn in purified CD14+ monocytes and following differentiation into moDCs (**Figure 5B-C**). Differentiation induced a profound remodeling of the FcR landscape, characterized by a marked reduction of CD64 expression and decreased levels of CD32 and FcRn in moDCs compared with their monocytic precursors (**Figure 5B-C**). We next assessed the modulation of FcR expression by IFN-α2 and IFN-β. The optimal doses of both IFN-I were determined by dose–response analyses, selecting doses that induced maximal CD86 upregulation and elicited comparable responses for both IFN-I subtypes (**Supplementary Figure 3**). Neither IFN-α2 nor IFN-β significatively modulated the expression of the activating FcγR CD16A or the FcRn. In contrast, IFN-β led to a significant increase in CD64 expression as well as a reduction of CD32A expression, which was not observed upon IFN-α2 (**Figure 5D**). Together, these findings demonstrate that monocyte differentiation into moDCs is accompanied by substantial FcR remodeling and reveal a selective effect of IFN-β, but not IFN-α2, in modulating FcγR expression levels, while maintaining the expression of FcRn. These results highlight distinct effects of IFN-I subtypes on FcR regulation in moDC.

**Figure 5:**
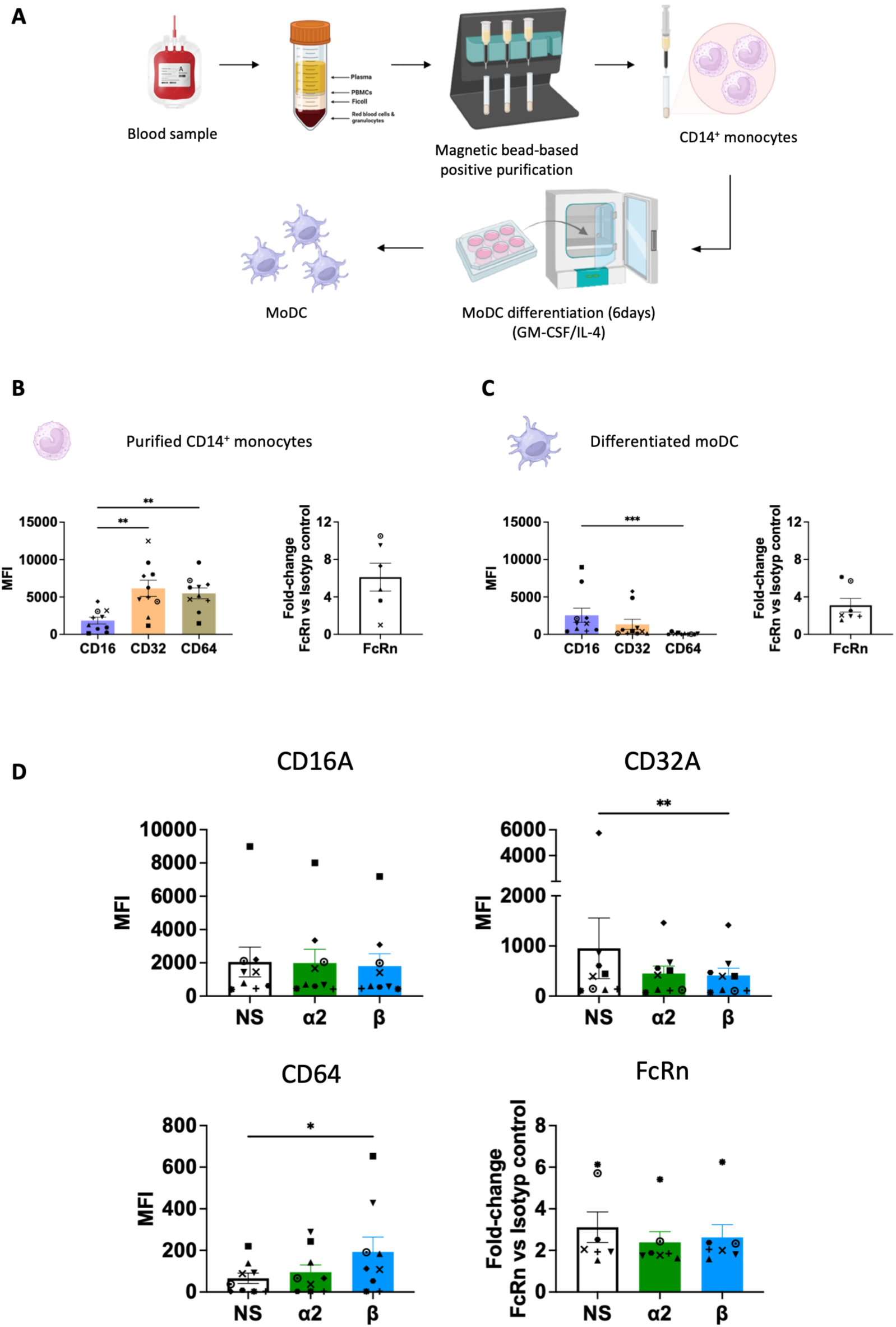
Workflow of generation of moDCs and their IFN-mediated FcR modulation. (A) Freshly isolated PBMCs were used as starting material for the positive selection of CD14^+^ monocytes. They were differentiated into moDC by culturing them with GM-CSF and IL-4 during 6 days. (B-C) Basal expression of FcRs in freshly purified CD14^+^ monocytes (B) and differentiated moDC (C), assessed by conventional cytometry. (D) Modulation of FcɣR (CD16, CD32, CD64) and FcRn expression, measured after stimulation of moDC by IFN-I (D). FcRn expression corresponds to the ratio of FcRn MFI and isotype control. Values are mean +/- SD of 9 for FcγR and 7 for FcRn, independent measurements. The concentrations tested are 1000 IU/mL for IFN-α2 and 100 pM for IFN-β (n>7). Statistics were performed by Wilcoxon test (* p ≤ 0.05, ** p ≤ 0.01, *** p ≤ 0.001).

#### IFN-β leads to enhanced phenotypic and functional activation of moDC as compared to IFN-α2

We next assessed the phenotypic and functional activation of moDC by IFN-α2 and IFN-β. We measured the modulation of cell surface markers associated with T-cell co-stimulation and antigen presentation (CD40, CD80, CD86, HLA-ABC, and HLA-DR), DC maturation (CD83), adhesion and migration (CD44, CD62L, and CD11c), as well as chemotaxis-associated receptors (CXCR4 and CCR5) (**Figure 6A**). Both IFN-I subtypes significantly increased HLA-ABC, CD40, CD80 and CD86 expression on moDC (**Figure 6A; Supplementary Figure 3**). However, IFN-β induced a broader maturation phenotype, characterized by significant upregulation of HLA-DR, CD86 and CD83, not observed upon IFN-α2 activation. Neither IFN-β nor IFN-α2 affected CD11c, CD62L, CXCR4 and CCR5 expression (**Figure 6A**). In parallel, we evaluated the functional consequences of IFN-I stimulation by analyzing the cytokine and chemokine secretion profile of treated moDCs (**Figure 6B**). Consistent with the phenotypic activation, IFN-β induced a broader and stronger secretory response compared with IFN-α2. Specifically, IFN-β significantly enhanced the production of IL-6, IL-10, IFN-λ1, CCL2, CCL5, CXCL9, CXCL10, and CXCL11, whereas IFN-α2 significantly increased only CXCL10 secretion (**Figure 6C**). Overall, these results provide a comparative and comprehensive study of phenotypic and functional activation of moDC by IFN-I subtypes. These findings demonstrate that both IFN-I subtypes promote moDC activation, but IFN-β induces a more pronounced maturation and functional activation program, characterized by enhanced expression of antigen-presentation and maturation markers as well as a broader cytokine and chemokine response.

**Figure 6.**
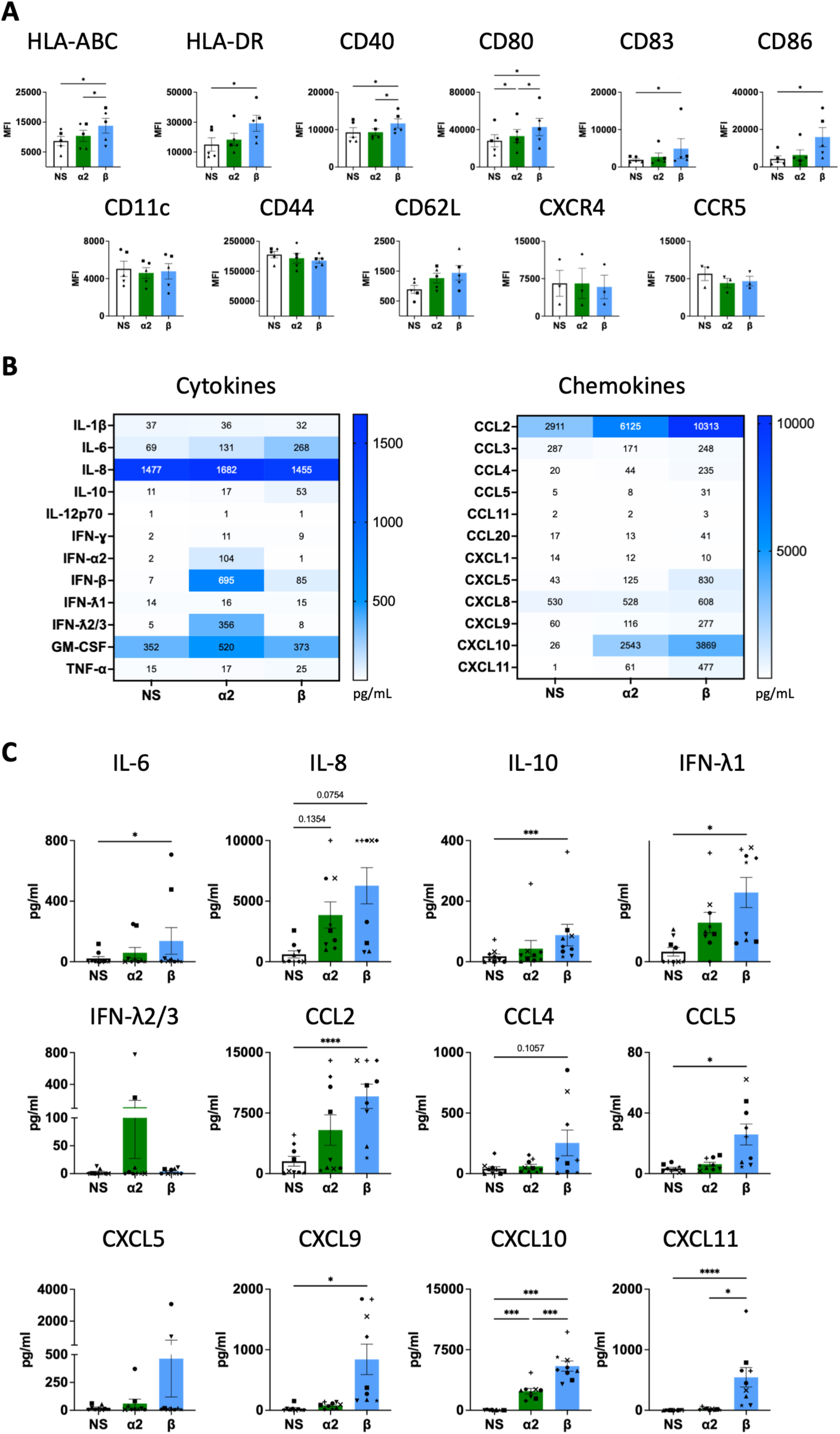
Phenotypic and functional activation of moDC after IFN-I stimulation. MoDc were generated as indicated in Figure 5A, and stimulated with IFN-I. (A) Phenotypic activation of moDCs assessed by monitoring cell surface markers using spectral flow cytometry. (B-C) Cytokine and chemokine secretion profile of IFN-I activated moDCs, represented as a heatmap (B) and histograms (C). The secretion of cytokines and chemokines was measured in pg/mL by LegendPlex assay. Values are mean +/- SD of 5 (A) and 9 (B-C), independent measurements (different symbols represent data from different HD samples). The concentrations tested are 1000 IU/mL for IFN-α2 and 100 pM for IFN-β (n > 7). Statistics were performed by Wilcoxon test or paired t test depending on the distribution normality (* p ≤ 0.05, ** p ≤ 0.01, *** p ≤ 0.001) (n ≤ 4).

### TAK-981 closely mirrors the global secretion profile induced by IFN-β in moDCs, while only partially recapitulating its effects on moDC phenotypic activation

We next assessed the effect of TAK-981 on moDC. Similar to IFN-β, TAK-981 modulated FcR expression by decreasing CD32A and increasing CD64 levels (**Figure 7A**), indicating that IFN-I induction through SUMOylation inhibition recapitulates the IFN-β-driven remodeling of the FcR profile in moDCs. Like IFN-β, TAK-981 also enhanced the expression of CD86, CD80 and HLA-ABC. However, in contrast to IFN-β, TAK-981 did not significantly increase HLA-DR or CD40 expression. Instead, TAK-981 reduced the expression of CD11c and CD44, changes that were not observed following stimulation with either IFN-β or IFN-α2, highlighting specific differences between pharmacological IFN-I induction and direct IFN-β stimulation (**Figure 7B**). Additionally, TAK-981 significantly increased the secretion of IL-8, IL-10, IFNλ1, TNFα, CCL2, CCL4, CCL5, CXCL9, CXCL10 and CXCL11 by MoDC (**Figure 7C-D**), resulting in a cytokine and chemokine profile that largely resembled that induced by IFN-β. Together, these findings demonstrate that TAK-981 recapitulates major aspects of the IFN-β-driven phenotypic and functional activation program in moDCs, including FcR remodeling and inflammatory mediator production, while inducing distinct changes in selected surface markers associated with moDC phenotype.

**Figure 7.**
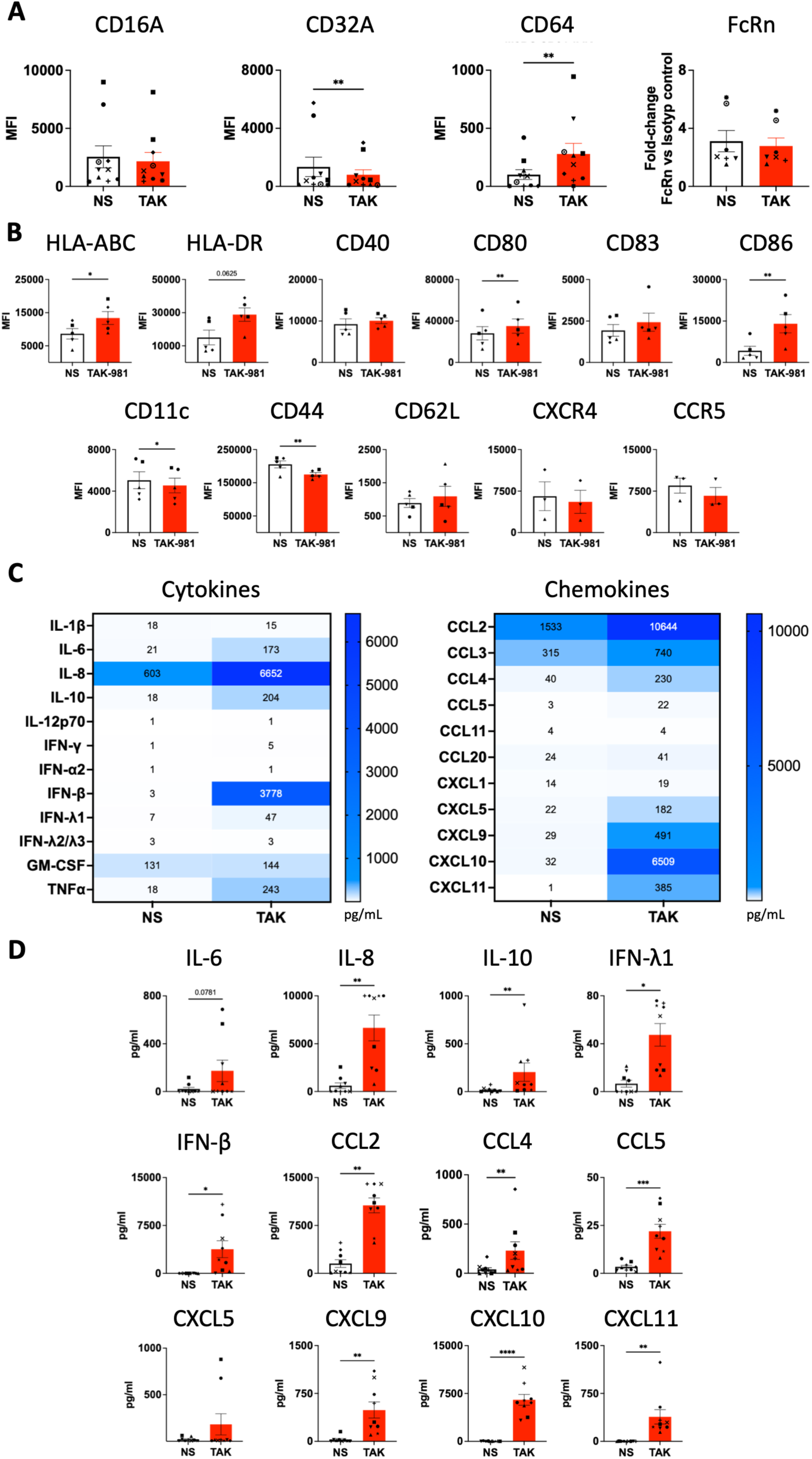
FcRs modulation and activation profile by moDC after TAK-981 stimulation. Modulation of FcɣR (CD16, CD32, CD64) and FcRn expression was observed after stimulation of moDC by TAK-981 at 100 nM (A). FcR expression on moDC, assessed by conventional flow cytometry. FcRn expression corresponds to the ratio of FcRn MFI and isotype control. (B) Phenotypic activation of moDC assessed by monitoring cell surface markers using spectral flow cytometry. (C-D) Cytokine and chemokine secretion profile of whole monocyte populations, represented as a heatmap (C) and histograms (D). The secretion of cytokines and chemokines was measured in pg/mL by LegendPlex. Statistics were performed by Wilcoxon test or paired t test depending on the distribution normality (* p ≤ 0.05, ** p ≤ 0.01, *** p ≤ 0.001) (n ≤ 4).

### IFN-I signaling induces redistribution of CD16A and spatial reorganization of Fc receptor pairs in moDCs

We next investigated whether IFN-I stimulation or TAK-981 treatment alters the spatial organization of FcR in moDCs, using the same ImageStream® approach applied to monocytes (**Supplementary Figure 2**). This analysis enabled the simultaneous assessment of FcR expression and spatial distribution following IFN-α2, IFN-β, or TAK-981 treatment. In moDCs, FcRn displayed a diffuse distribution, whereas CD16A was predominantly distributed in a circumferential pattern under basal conditions. Following IFN-I pathway activation or TAK-981 treatment, CD16A adopted a more dispersed, punctate distribution throughout the cell. Analysis of receptor correlation using the Similarity Index revealed that most FcR pairs displayed limited spatial association, with the exception of FcRn and CD16A (**Figure 8A**). We next quantified stimulus-induced changes in FcR spatial organization by calculating the fold change in receptor relocalization index. This analysis identified selective alterations in FcR proximity following IFN-I pathway activation. In particular, IFN-I pathway activation and TAK-981 treatment promoted selective changes in FcR spatial organization, characterized by increased proximity between FcRn and CD32A, as well as between CD32A and CD16A, the latter representing the predominant relocalization pattern (**Figure 8B**).

**Figure 8.**
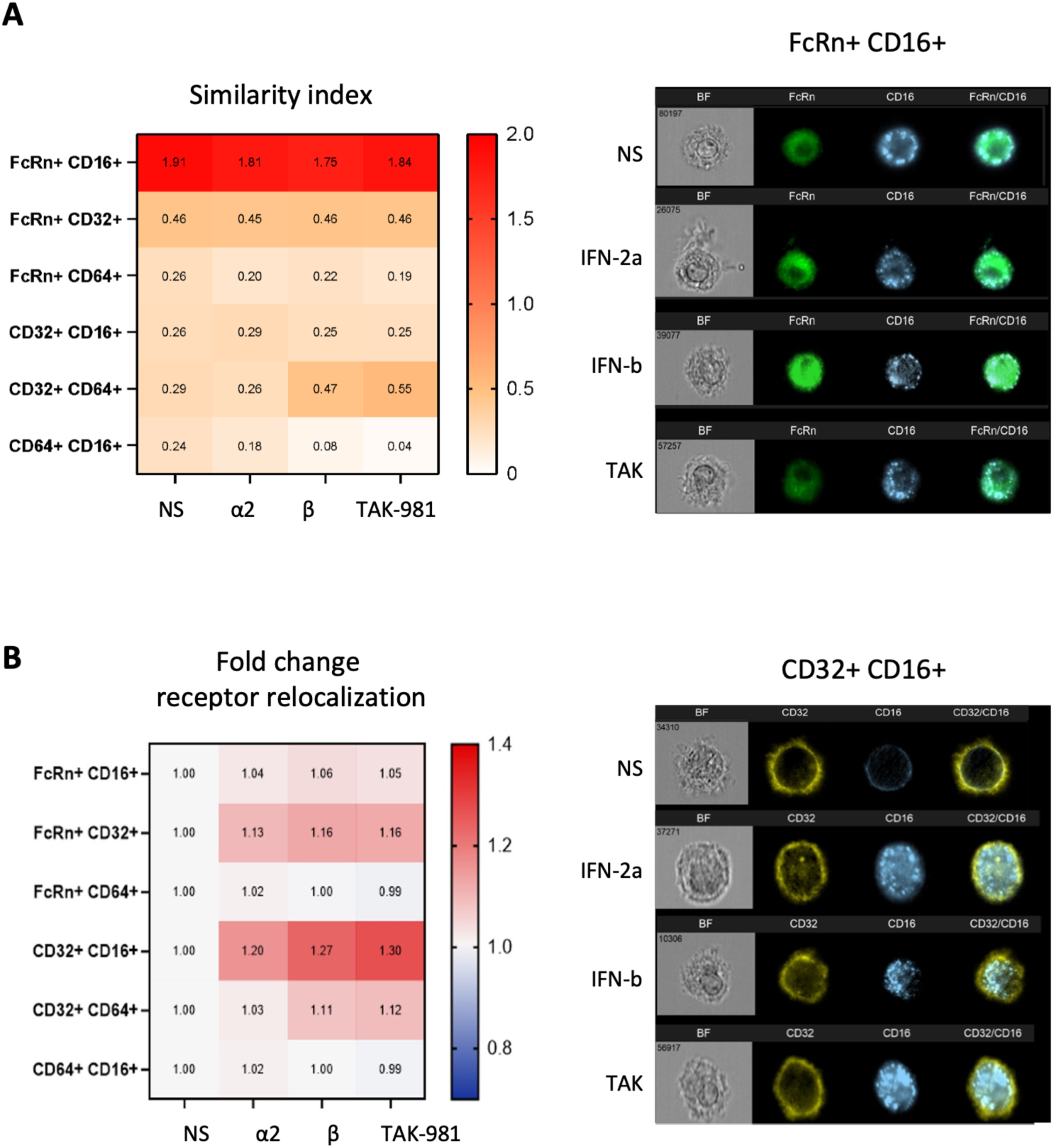
ImageStream analysis of FcR distribution and colocalization under various stimulations. MoDCs were generated from CD14⁺ monocytes isolated from the peripheral blood of healthy donors by magnetic bead-based positive selection and stimulated with IFN-α2 at 1000 IU/mL, IFN-β at 100 pM or TAK-981 at 100 nM. (A) Similarity index heatmap for the different receptors, generated using IDEAS® software, enabling the compartmentalization of membrane and intracellular regions. Fluorescence images showing respective receptor expression and their merge after stimulation for the receptor pair with the highest level of co-expression (n = 2). (B) Fold change heatmap of Bright Detail Similarity R3 (BDS) scores calculated using the IDEAS® automated wizard to quantify colocalised receptor expression. Images are representative of two independent donors.

Together, these findings suggest that IFN-I signaling and pharmacological IFN-I induction influence not only Fc receptor expression but also their spatial organization at the surface of moDCs, revealing an additional layer of Fc receptor regulation during myeloid cell activation.

## DISCUSSION

We have previously demonstrated in a mouse model of retroviral infection that both IFN-I and the inflammatory environment associated with retroviral infection enhance the expression of activating FcγRs ^26^. Here, we provide new insights into differential effects of IFN-α2, IFN-β and the IFN-I inducer, TAK-981, on FcR modulation in human monocyte subsets and moDCs. Our study also characterizes the secretome and immune receptor expression profiles of these distinct myeloid cell populations in response to different types of IFN-I, revealing stimulus-specific activation signatures. In addition, we provide a detailed and comprehensive study of the direct effects of TAK-981 on human myeloid cells. This has not been reported thus far, as previous studies have primarily focused on the activation of the mouse or human myeloid compartment as a whole, rather than isolated primary cells ^41,44^. This is an important point, as TAK-981 may indirectly activate immune cells by inducing the release of soluble mediators from neighboring cells that are directly responsive to the compound^41^. Overall, our research provides new understanding of how distinct IFN-I/IFN-I inducers differentially regulate the expression and cellular distribution of FcR in monocyte subsets and moDCs, and shape their cytokine/chemokine secretion profile (**Figure 9**).

**Figure 9.**
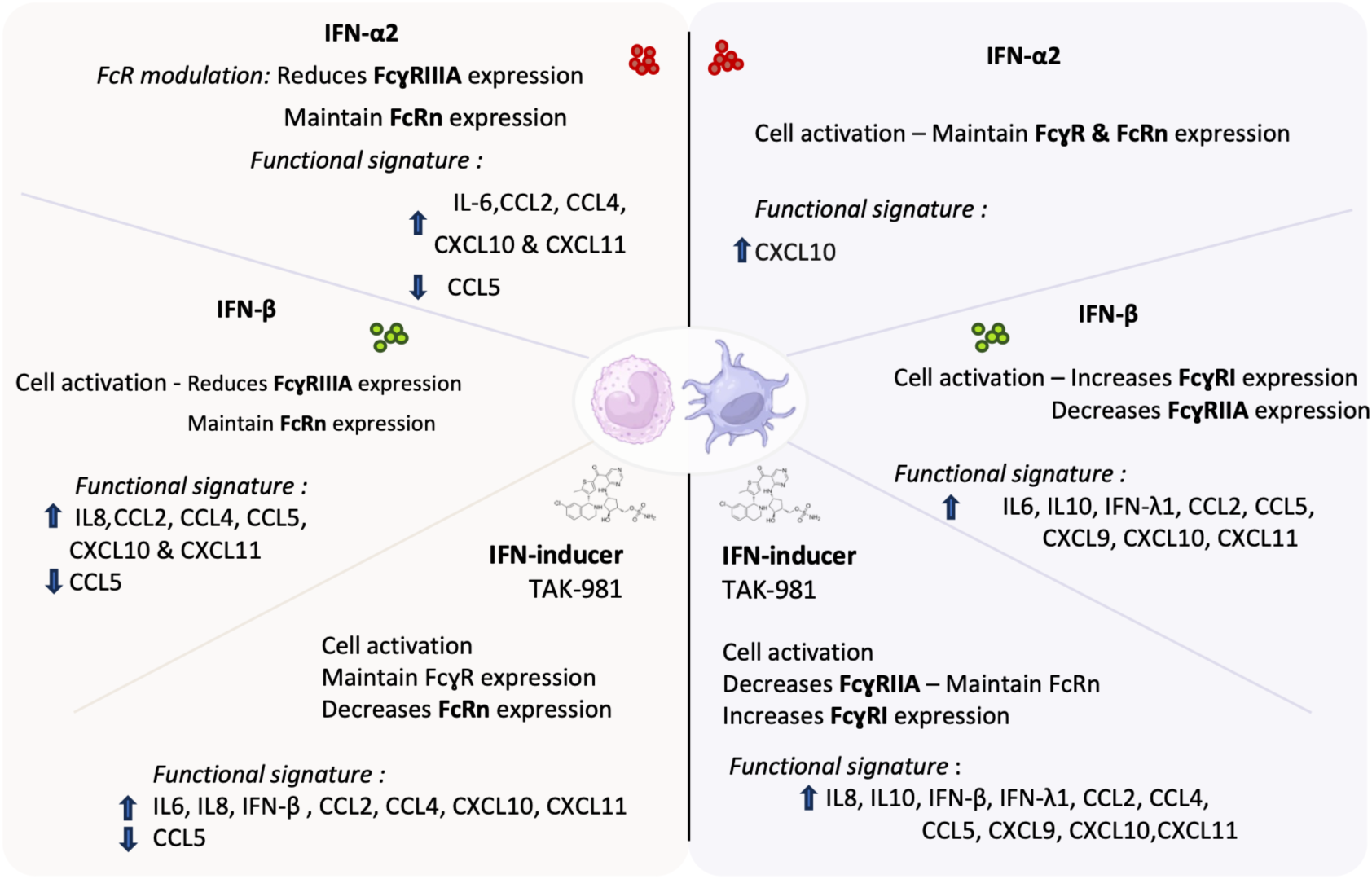
Overview of monocytes and moDC stimulation profiles by IFN-I/IFN-inducer stimulations. Summary of cellular activation, FcRs modulation and secretion profiles observed after monocytes or moDC stimulation by IFN-I (IFN-α2 and IFN-β) or the interferon-inducer TAK-981.

The effect of different IFN-I subtypes on FcR expression in monocyte subsets remains poorly understood. This is especially true for FcRn, as existing studies are limited and primarily focus FcRn gene expression in THP-1 cells rather than primary monocytes ^50^. We found that IFN-α2 and IFN-β decrease the cell surface levels of CD16A particularly in classical monocytes, but not in the NC subset. This observation aligns with recent reports describing CD16A downregulation upon IFN-α-mediated activation of monocytes ^49,51^. However, these studies primarily relied on gene expression analyses of IFN-α-activated PBMCs rather than isolated primary monocytes, making it difficult to distinguish direct effects of IFN-I on monocyte subsets from indirect effects mediated by other IFN-I-activated cells. Additionally, these studies did not specify monocyte subsets or IFN-α subtypes, nor did they assess the effects of IFN-β. Interestingly, CD16A downregulation has also been observed in human NK cells activated by IFN-I inducers ^52^. Our findings contrast with previous results in mouse monocytes, where IFN-I stimulation enhanced the expression of activating FcγRs ^26,53^ highlighting species-specific differences in IFN-I-mediated FcR modulation. Worthy of note, neither IFN-α2 nor IFN-β modulated the expression levels of FcRn. This is an important novel finding as modulation of FcRn has mainly been reported upon IFN-γ or TNF-α activation of THP-1 cells ^50,54^. The maintenance of FcRn levels despite extensive IFN-I-driven activation suggests that IFN-I selectively regulate immune effector pathways while preserving mechanisms required for IgG homeostasis. This finding has important implications for antibody-based therapies, as FcRn represents a central determinant of IgG recycling, serum persistence, and tissue distribution. In particular, therapeutic antibodies incorporating Fc modifications designed to enhance FcRn interaction, such as LS- or YTE-engineered antibodies, may be influenced by inflammatory environments that alter FcγR expression while maintaining FcRn-dependent pharmacokinetic properties. Similarly to monocytes, we found that IFN-α2 and IFN-β also modulated the expression levels of activating FcγRs in moDC, in particular CD32A instead of CD16A, while not affecting the levels of FcRn. In contrast, IFN-β but not IFN-α2, enhances the expression of the high affinity FcγR CD64. Interestingly, previous studies by Dugast *et al.* in the context of HIV-1 infection have reported differential modulation of FcγR expression in myeloid cells during acute versus chronic infection ^55^. Specifically, acute infection was associated with increased CD64 expression, whereas chronic infection was characterized by decreased CD32 and CD16 expression across several myeloid cell subsets, including mDCs, pDCs, and monocytes. These changes in FcγR expression might be consistent with the effects of IFN-I on FcγR modulation that we observed, suggesting that the changes in FcγR expression during HIV-1 infection may reflect an evolving IFN signature over the course of infection. Noteworthy, the increased expression of CD64 was also observed in TAK-981 activated moDC, in agreement with the production of IFN-β by cells activated by this IFN-inducer. However, we observed some differences in FcR modulation between monocytes activated by IFN-I and those activated by TAK-981. Thus TAK-981 maintained the expression level of all FcγRs on monocytes but slightly downregulated FcRn in classical monocytes. Taken together, our findings demonstrate a type-specific and cell-specific effect of IFN-I on FcR modulation which might need to be taken into account in combinatorial therapeutic approaches using mAb and IFN-I/IFN-inducers. This may be particularly relevant for Fc-engineered therapeutic antibodies designed to enhance binding to specific FcγRs or FcRn. The development of such Fc-engineered antibodies is an expanding field aimed at improving the efficacy and therapeutic potential of antibody-based therapies^21^. Thus, understanding how cytokine environments reshape FcR expression and function may become increasingly important for the rational design and optimization of advanced antibody therapeutics

Beyond changes in receptor abundance, our imaging analyses suggest that IFN-I may regulate FcR receptor organization at the cell surface. In monocytes, CD16A predominantly remained polarized toward one pole of the cell under basal conditions and following TAK-981 treatment, whereas IFN-α2 and IFN-β promoted a more dispersed, punctate distribution throughout the cell. In moDCs, CD16A displayed a circumferential distribution under basal conditions but adopted a more dispersed, punctate pattern following IFN-I pathway activation. Thus, IFN-I signaling was associated with distinct changes in FcR distribution patterns, with more pronounced spatial redistribution observed in moDCs. These observations suggest that IFN-I signaling and pharmacological IFN-I induction regulate not only the abundance of FcRs but also their cellular positioning and distribution patterns, revealing an additional layer of FcR regulation that differs according to both cell type and stimulus.

Immune receptor signaling is increasingly recognized to depend not only on expression levels but also on spatial arrangement, nanoscale clustering, membrane mobility, and interactions with cytoskeletal structures. Although the functional consequences of the FcR redistribution observed here remain to be determined, these findings raise the possibility that IFN-I influence antibody-mediated responses by altering both the quantity and the organization of available receptors. Such spatial regulation could theoretically modify receptor accessibility, signaling thresholds, or coordination with intracellular signaling machinery. Future studies combining super-resolution imaging, biochemical approaches, and functional assays will be required to determine whether IFN-I-induced FcR reorganization directly influences processes such as phagocytosis, antibody-ADCC, or immune complex signaling.

Our findings also provide insights into the differential phenotypic and functional activation of monocytes and moDC upon activation by IFN-α2 and IFN-β. Both IFN-I types lead to an increase in the expression of CD86 and CD62L as well as a decrease of the chemokine receptor CXCR4 on both monocyte subsets. In addition, IFN-β but not IFN-α2 also lead to reduced cell levels of CCR5 in classical monocytes. On the contrary, CXCR4 and CCR5 were not modulated by IFN-I subtypes in moDC. These observations expand current knowledge of the effect IFN-I on chemokine receptor expression from studies describing the effects of IFN-α2 on the increase of CCR1 and CCR3 in primary mononuclear phagocytes, as well as in the monocytoid cell line U937^56^. Worthy of note, IFN-β led to a stronger phenotypic activation of moDC as compared to IFN-α2 revealed by the enhanced expression of both MHC molecules (HLA-DR et HLA-ABC) as well as co-stimulatory molecules (CD40, CD80, CD86 and CD83) not observed upon IFN-α2 activation. Our findings are in agreement with previous studies^57^, notably the higher expression of CD80, CD86 and HLA-ABC observed by IFN-β versus IFN-α2 activation. The higher phenotypic activation of moDC upon stimulation by IFN-β was also associated with an enhanced secretion profile characterized by the production of IL-6, IL-10, IFNλ1, CCL2, CCL5, CXCL9, CXCL10 and CXCL11, at difference of IFN-α2 that only significatively enhanced CXCL10, providing clear evidence for stronger activation of moDC by IFN-β, both qualitatively (types of chemokines/cytokines secretion) than quantitatively (secretion level/amount). These results are consistent with previous findings by Garcin *et al.*, who reported a more pronounced chemokine secretion profile in IFN-β– versus IFN-α2–differentiated moDCs, despite important differences in the experimental approaches used. Specifically, whereas Garcin *et al.*^58^ induced monocyte differentiation into moDCs using IL-4 in combination with IFN-I subtypes, we previously differentiated monocytes into moDCs using IL-4 and GM-CSF, followed by activation with distinct IFN-I subtypes. In contrast to these observations, both IFN-I types led to a similar secretion profile in activated monocytes, characterized by increased secretion of CCL2, CCL4, CXCL10 and CXCL11, but the differences were mostly quantitative, with stronger secretion by IFN-β-activated monocytes. These observations align with previous reports on monocyte-derived macrophages, which describe an IFN-β-driven “beta-chemokine” program that promotes the recruitment of monocytes, T cells, and NK cells ^59^. Thus, both IFN-α and IFN-β drive human monocytes toward a chemokine profile that favors mononuclear cell recruitment and Th1 antiviral immunity^60^, although to a different extent.

Our study extends the knowledge on the immunomodulatory properties of TAK-981 by fine dissecting its effect on the phenotypic and functional activation of monocyte subsets and moDC. It provides additional insight into how pharmacological induction of endogenous IFN-I responses differs from direct cytokine stimulation. This novel IFN-I inducer has been show to activate several innate immune cells and enhances the efficacy of anti-cancer therapeutic antibodies, notably in preclinical mouse studies ^43^. However, its effects on human myeloid cells remain poorly characterized, as the few studies reported rely on global PBMC analyses rather than isolated primary cells, making it difficult to distinguish direct effects on specific cell types from secondary cytokine-mediated activation ^41,45^. By examining purified human myeloid populations, we show that TAK-981 largely reproduces the IFN-β-driven activation profile, consistent with its ability to induce endogenous IFN-I production. Nevertheless, TAK-981 effect was not identical to that of recombinant IFN-β, displaying cell-specific differences in the modulation of selected FcR, cell surface activation markers and cytokine/chemokine responses, in monocytes versus moDC. These differences likely reflect the complexity of endogenous IFN induction, which may involve distinct kinetics, autocrine amplification loops, and additional inflammatory mediators generated downstream of pathway activation. Thus, TAK-981 should not simply be considered an IFN-β substitute, but rather as a pharmacological strategy capable of establishing an IFN-associated immune state. Thus, when intended for use as an adjuvant or immunomodulatory agent, these differential effects should be considered based on the targeted therapeutic outcome.

Overall, this study provides a comprehensive and comparative analysis of IFN-I subtypes and the IFN-I inducer TAK-981, integrating multiple phenotypic and functional parameters of myeloid cell activation (**Figure 9**). Our findings reveal that IFN-I regulate human myeloid immunity through multilayered remodeling of the FcR landscape—including changes in expression levels and spatial organization—as well as cellular activation and secretory programs. The magnitude and nature of these effects depend on the IFN-I subtype, with IFN-β eliciting the most extensive response, while TAK-981 recapitulates many, but not all, features of the IFN-β-induced state. The biological and therapeutic implications of these findings extend beyond IFN-I biology. During viral infections, IFN-I-mediated remodeling of FcRs may influence antibody effector functions, including vaccine-induced antibody responses and the activity of therapeutic mAbs whose efficacy depends on FcγR engagement. This may be particularly relevant for Fc-engineered mAbs designed to selectively enhance interactions with activating FcγRs or FcRn. Furthermore, the ability of TAK-981 to induce an IFN-associated myeloid state supports its continued investigation as an IFN-I-inducing strategy for combination with antibody-based immunotherapies. Taken together, our findings may inform the rational design of vaccine and therapeutic strategies that harness IFN-I signaling, particularly in combination with Fc-engineered antibodies, where cytokine-driven changes in FcR expression and organization may influence antibody activity.

## MATERIALS AND METHODS

### Ethics

Human blood samples were obtained from healthy adult anonymous donors at the regional blood bank (Etablissement Français du sang, EFS, Montpellier, France). The study was approved by the Occitanie & Midi-Pyrénées EFS scientific board (EFS-OCPM: N° PLER2024-007-R). All donors provided written informed consent.

### Primary cell purification

Peripheral blood mononuclear cells (PBMC) were isolated from buffy coat on a Ficoll-Histopaque® 1077 gradient (Sigma-Aldrich, Lyon, France).

For the study of untouched monocytes, cells were purified from PBMCs using Pan Monocyte Isolation Kit, human (Miltenyi, Cat#130-096-537) according to the manufacturer’s protocol. Purity assessed by flow cytometry was >95%

For moDC generation, monocytes were isolated from PBMCs by CD14^+^ expression by positive immunomagnetic selection (MACS system, MiltenyiBiotec, cat#130-050-201) as previously described in Paris et al ^61^. CD14^+^ cells were incubated for 6 days at 2 x 10^6^ cells per well, in the 6-wells plate, in the presence of 50 ng/ml granulocyte-macrophage colony-stimulating factor (GM-CSF) and 20 ng/ml interleukin-4 (IL-4) (BioTechne R&D System, Abingdon, United Kingdom, cat#7954-GM-050/CF & cat#6507-IL-025/CF). At day 3, 1 mL of complete media was added in each well. The medium used for culture was RPMI 1640 supplemented with GlutaMAX™ (Gibco, cat#61870-010), 10% Foetal Bovin Serum (FBS, Eurobio, REF#CVFSVF00-01) and 1% Penicillin/Streptomycin (Pen Strep Gibco, cat #15140-122). At day 6, moDCs were ready for stimulation tests.

### Cell culture

Purified monocytes were plated at a concentration of 1 million cells/ml in 150 μl in 96 U-bottom wells and moDCs were plated at a concentration of 6 x 10^5^ cells/mL in 300µL (2.10^6^ cells/mL) in 24-wells plates and cultured in RPMI 1640 supplemented with GlutaMAX™ (Gibco, cat#61870-010), 10% FBS (Eurobio, REF#CVFSVF00-01) and 1% Penicillin/Streptomycin (Pen Strep Gibco, cat #15140-122). Cells were cultured for 18h or monocytes and 16h for moDCs with or without stimulation.

### In vitro stimulation

Monocytes were stimulated for 18h with IFN-α2a (Pbl assay science, Cat#11101-1) at 10 000 U/ml, IFN-β (Biogen, Avonex, Idec Inc Cambridge, MA, USA) at 1 nM, or TAK-981 (Subasumstat, MedChemExpress, Cat#HY-111789) at 100 nM. MoDCs plated at 6 x 10^5^ cells in 300 µL of the complete medium described above, stimulated for 16h with IFN-α2a (Pbl assay science, Cat#11101-1) at 1000 IU/ml, IFN-β (Biogen, Avonex, Idec Inc Cambridge, MA, USA) at 100 pM, or TAK-981 (Subasumstat, MedChemExpress, Cat#HY-111789) at 100 nM.

These doses were identified by dose-response experiments using different concentrations of IFN-α2a, IFN-β and TAK-981. The range tested were: IFN-α2 from 100 IU/mL to 10 000 IU/mL (equal to 0,5 nM, 5,2nM and 52 nM, respectively); IFN-β from 1 pM to 500 pM; Subasumstat (TAK-981) at 100 nM and 1 µM.

### Flow cytometry

Monitoring of cell phenotypes was done by flow cytometry using FACSCantoII (BD Bioscience) and Cytek Aurora (Cytek Biosciences) flow cytometers and analysis was performed using Flowjo software 10.10.

Purified monocytes were first stained with a viability dye (Zombie NIRTM, Biolegend #423105), washed and stained with fluorochrome-conjugated antibody mix for 15 min at room temperature. Antibodies for flow cytometry were CD14-PerCp-Cy5.5 (M5E2, BD Pharmingen, Cat #550787), CD16-PE-Cy7 (3G8, Biolegend, Cat #302015), CD32-PE (FUN-2, Biolegend, Cat #303206), CD64-APC (10.1, Biolegend, Cat #305014), CD86-BV421 (BU63, Biolegend, Cat #374212), HLA-DR-BV510 (L243, Biolegend, Cat #307646), CD62L-BV605 (DREG-52, Biolegend, Cat #304834), CD184-BV785 (12G5, Biolegend, Cat #306529), CD195-BV711 (J418F1, Biolegend, Cat #359129), CD317-AF647 (RS38E, Biolegend, Cat #348404), hFcRn-AF488 (937508, R&D System, Cat #IC8639G). For FcRn intracellular detection, cells were washed and then fixed and permeabilised using the Cytofix/Cytoperm kit (BD, Cat #554714) during 30min à 4°C, dark. Cells were incubated with the hFcRn-AF488 antibody for 20 min at 4°C, dark. A mouse IgG2b Alexa Fluor 488 antibody (clone 133303, Cat. #IC0041G, R&D Systems) was used as the isotype control for FcRn staining and to determine the fold change in FcRn expression. Same conditions of intracellular staining were used for moDC.

Before the differentiation into moDC, CD14^+^ monocytes were phenotyped for CD14 PE (M5E2, cat#555398, BD Pharmingen™), CD32 PE (FUN-2, cat#303206, BioLegend®), CD16 PE-Cy7 (3G8, cat#302016, BioLegend®), CD64 APC (10.1, cat#305014, BioLegend®) and FcRn AlexaFluor 488 (937508, cat#IC8639G, R&D System) for donors analyzed with both CANTO II (conventional cytometry). A mouse IgG2b AlexaFluor 488 (Clone 133303, CAT IC0041G, R&D System) isotypic control was used to determine the fold change for FcRn.

To assess the phenotypic properties of moDCs following different stimulation conditions and to characterize FcR modulation, a six-color panel was used for samples analyzed on the FACSCanto II. The panel included CD11c-PerCP-Cy5.5 (3.9, Cat. #301623, BioLegend®), CD16-PE-Cy7 (3G8, Cat. #302016, BioLegend®), CD32-PE (FUN-2, Cat. #303206, BioLegend®), CD64-APC (10.1, Cat. #305014, BioLegend®), and FcRn-Alexa Fluor 488 (937508, Cat. #IC8639G, R&D Systems). The mouse IgG2b Alexa Fluor 488 isotype control (clone 133303, Cat. #IC0041G, R&D Systems) was used as the control for FcRn intracellular staining. A second panel was then conceived for a broader immunophenotyping of activated moDC to be assessed using the Cytek Aurora cytometer. Antibody for activation profile were CD11c PerCP-Cy5.5 (3.9, cat#301623, BioLegend®), CD40 PE-Dazzle (5C3, 334341, BioLegend®), CD44 AlexaFluor 700 (BJ18, cat#338813, BioLegend®), CD62L BV605 (DREG-52, cat#304834, BioLegend®), CD80 RB780 (2D10.4, cat#755432, BD OptiBuild), CD83 BV650 (HB15e, cat#740602, BD OptiBuild), CD86 BV421 (BU63, cat#374212, BioLegend®), HLA-ABC PerCP (W6/32, cat#311421, BioLegend®) and HLA-DR BV510 (L243, cat#307647, BioLegend®). To observe the modulation of FcRs, antibodies were CD16 PE-Cy7 (3G8, cat#302016, BioLegend®), CD32 PE (FUN-2, cat#303206, BioLegend®), CD64 APC (10.1, cat#305014, BioLegend®) and FcRn AlexaFluor 488 (937508, cat#IC8639G, R&D System). The monitoring of cellular receptor was with CCR5 BV711 (J418F1, cat#359129, BioLegend®) and CXCR4 BV785 (12G5, cat#306529, BioLegend®). Antibody volumes were adapted to the number of cells based to the manufacturer’s instructions. Cells were incubated for 25 min at 4°C with agitation. The cells were then washed twice (400xg, 4°C, 5 min) and resuspended in 150 uL of buffer (PBS). The 7-AAD (7-aminoactinomycin D) marker (cat#420404, Biolegend®) is added at 1/250th 10 min before reading to observe cell viability in each sample for both cytometry system. Forward scatter area and forward scatter height as well as side scatter area and height were used to remove doublets from flow cytometry analyses. Gating is done on live cells as assessed by viability dye and on moDCs. Data were analyzed using FlowJo software version 10.10.0 (FlowJo LLC 2025).

### Soluble cytokines and chemokines quantification

Monocytes and moDC were activated by the different stimuli for 18h for monocytes and 16h for moDCs. Supernatants were collected and soluble cytokines/chemokines were measured using two panels of bead-based immunoassays (Cytokine Bead Array) following the manufacturer’s instructions. The following panels were used: (i) the LEGENDplex™ HU Anti-Virus Response Panel (13-plex, cat#741270, Biolegend®), allowing the quantification of IL-1β, Il-6, TNF-α, IP-10, IFN-λ1, IL-8, IL-12p70, IFN-α2, IFN-λ2/3, GM-CSF, IFN-β, IL-10, IFN-ɣ and (ii) the LEGENDplex™ HU Proinflammatory Chemokine Panel (13-plex, cat#740985, Biolegend®), allowing the quantification of IL-8, CXCL10, CCL11, CCL17, CCL2, CCL5, CCL3, CXCL9, CXCL5, CCL20, CXCL1, CXCL11, CCL4. The concentration of each analyte was quantified by Novocyte (Agilent Technologies) flow cytometer and data were analyzed with Biolegend LEGENDplex™ software provided by the manufacturer (https://legendplex.qognit.com).

### ImageStreamX MkII

For monocytes and moDC staining, a 5-panel immunophenotyping was done with ImageStreamX MkII. To observe the modulation and location of FcRs, antibodies were CD16 PE-Cy7 (3G8, cat#302016, BioLegend®), CD32 PE (FUN-2, cat#303206, BioLegend®), CD64 APC (10.1, cat#305014, BioLegend®) and FcRn AlexaFluor 488 (937508, cat#IC8639G, R&D System). DAPI was used to stain the cell nucleus (cat#422801, Biolegend). The labeling procedure is performed in 1.5mL Eppendorf tubes at a cell concentration of 20M/mL with a minimum final volume of 25µL. Antibody volumes were adapted to the number of cells based to the manufacturer’s instructions and the sensitivity of the ImageStreamX lasers. Cells were incubated for 25 min at room temperature (RT) with agitation in the dark. The cells were then washed twice (400xg, RT, 5 min) and resuspended for the permeabilization step in 200 uL of BD Cytofix/Cytoperm™ Fixation/Permeabilization Kit (cat#554714, BD Biosciences) during 25 min at 4°C in the dark. The washing step is repeated in exactly the same way. Next, the cells were incubated with an anti-FcRn antibody or a control isotype antibody for 30 minutes in the dark at 4°C before being washed as before. DAPI labeling was added at 1µM during 5min at RT and washed once (400xg, RT, 5 min). Cells were resuspended for the acquisition at a concentration of 20M/mL. Doublets were removed (Area vs Aspect ratio from Brightfield channel) from the focused cells, during acquisition with ImageStreamX (INSPIRE software). Data were analyzed using IDEAS software. For each pair of receptors, we applied the “Similarity” feature to the membrane and cytolasm masks, and also used the “Colocalization” wizard with the “Bright Detail Similarity R3” feature.

### Statistical analysis

Statistical analysis was performed using Graph Pad Prism 10.6.1. Statistical comparisons were made with Shapiro-Wilk test to assess normality. Simple group comparisons were performed using Paired-t-test for parametric distribution and Wilcoxon for nonparametric distribution. Multiple group comparisons were performed using One-Way ANOVA with Geisser-Greenhousse correction for parametric distribution or Friedman test for nonparametric distribution. P-values of pairwise comparisons were adjusted for multiple comparisons using Tukey’s mulitple comparisons test for One-Way ANOVA and Dunn’s multiple comparison test for Friedman test. Significant differences were noted when p < 0.05. (* represents p < 0.05, ** represents p < 0.01 and *** represents p < 0.001).

## Supporting information

Supplementary Figures and references

## ACKNOWLEDGMENTS

This work was supported by grants from Sidaction (23-1-AEQ-13608, 2023-1-FJC-13614) and ANRS (France Recherche Nord&sud Sida-hiv Hépatites; ECTZ347813). Remi Bru, Océane Paris, M. Naranjo-Gomez and M. Pelegrin are members of the “MabImprove Labex”, a public grant overseen by the French National Research Agency (ANR) as part of the “Investments for the future” program (reference: ANR-10-LABX-53-01) that also supported this work. We thank the imaging facility MRI, which is part of the UAR BioCampus Montpellier and a member of the national infrastructure France-BioImaging, supported by the French National Research Agency (ANR-10-INBS-04, “Investments for the future”). We are grateful to Christophe Duperray et Felicia Leccia (MRI), for their help in implementing spectral cytometry procedures and to Quiterie Labaye for providing help in data analysis.

## AUTHOR CONTRIBUTIONS

Remi Bru (RB), Océane Paris (OP), Mar Naranjo-Gomez (MN-G) and Mireia Pelegrin (MP) defined the research program; RB, OP and Stephanie Boireau (SB) performed the experiments; Guillaume Bossis (GB) provided preliminary data on TAK-981 activation of monocytes. RB, OP, MN-G and MP carried out the data analyses. MN-G and MP wrote the manuscript. Grants to RB, OP and MP funded the study. All authors discussed, commented on and approved the manuscript in its final form.

