## Supplementary Figures and references for "Differential Effects of Distinct Type I Interferons and IFN Inducers on Fc Receptor landscape and myeloid cell function"

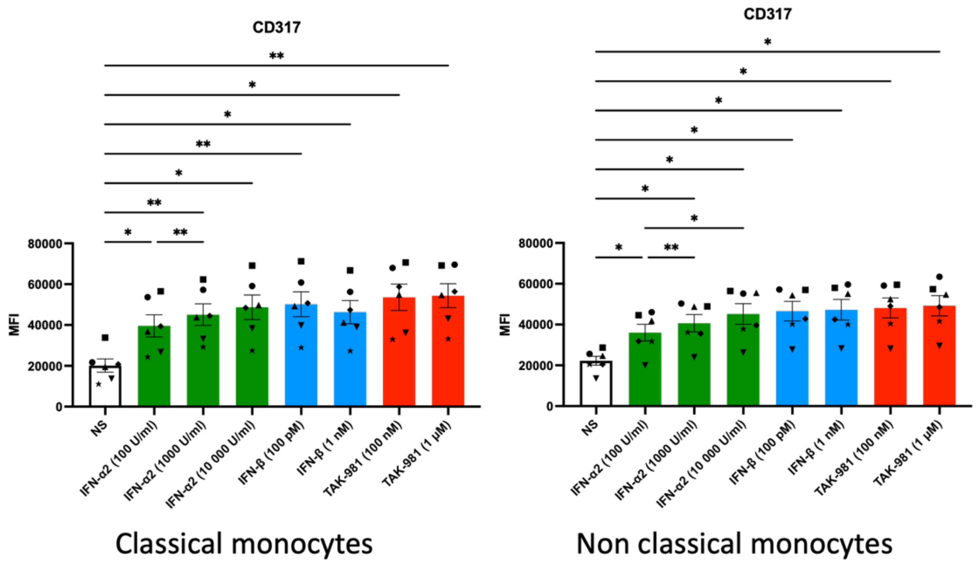


**Supplementary Figure 1. Dose-response experiments for identification of the optimal dose of IFN-α2, IFN-β and TAK-981.** Monocytes from healthy donors were purified from blood samples by magnetic bead-based negative selection and activated by increasing doses of IFN-α2, IFN-β and TAK-981. The doses that induced plateau levels of the type I IFN-inducible marker CD317 and elicited comparable responses for all compounds were selected for further experiments. Values are mean +/- SD of 6 independent measurements (different symbols represent data from different HD samples).

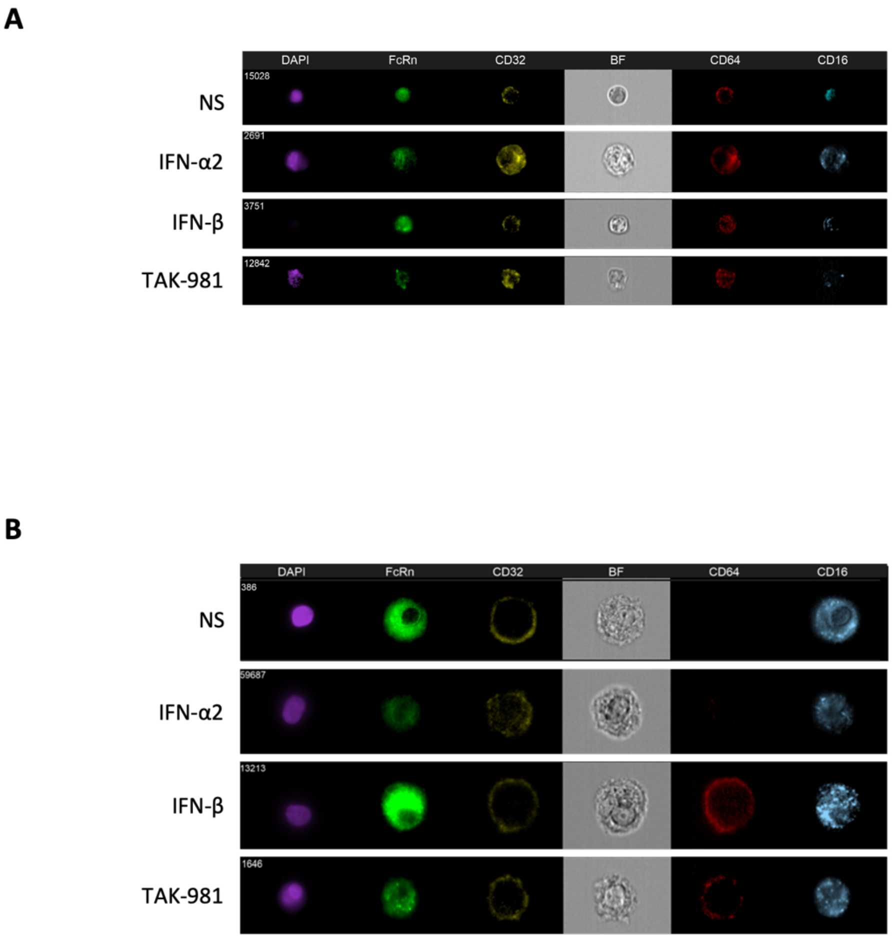


**Supplementary Figure 2. ImageStream® analysis of Fc receptor expression and spatial distribution in monocytes and moDCs following IFN-I pathway activation**. Representative ImageStream® images of (A) human monocytes and (B) monocyte-derived dendritic cells (moDCs) under basal conditions (NS) or following treatment with IFN-α2, IFN-β, or TAK-981. Cells were stained for FcRn, FcγRIIA (CD32A), FcγRI (CD64), and FcγRIIIa (CD16A), with nuclei visualized by DAPI. Brightfield (BF) images are shown to provide cellular morphology. The images illustrate stimulus-dependent differences in Fc receptor expression and spatial distribution, including changes in the localization and distribution pattern of CD16A following IFN-I pathway activation. Images are representative of n ≥ 2 donors.

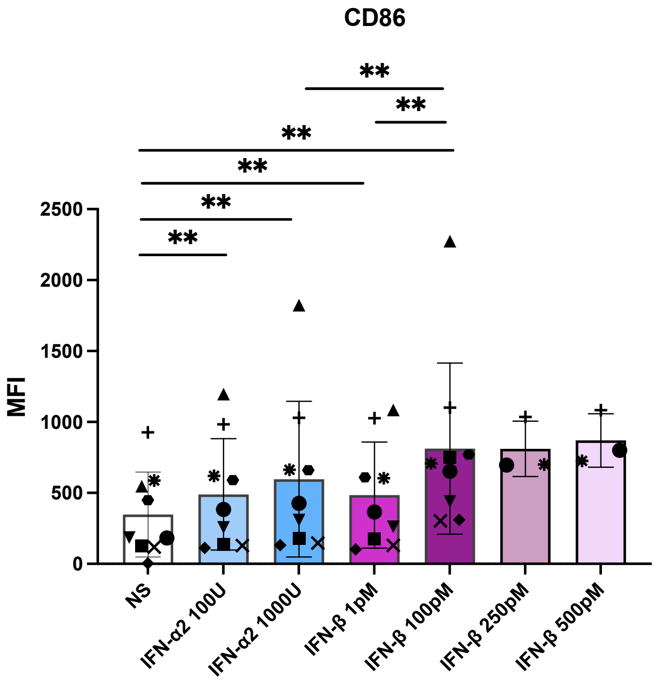


**Supplementary Figure 3. Dose-response experiments for identification of the optimal dose of IFN-α2 and IFN-β.** MoDCs were generated from CD14$^+$ monocytes isolated from the peripheral blood of healthy donors by magnetic bead-based positive selection and stimulated with IFN-α2 or IFN-β. Based on previous studies reporting robust moDCs activation at IFN-α2 concentrations ranging from 30 to 1000 IU/mL, 100 and 1 000 IU/mL were selected as biologically relevant doses and tested. Higher doses 10,000 IU/mL, were not included as they have not been shown to provide additional benefit in this experimental setting and avoid unnecessarily high IFN-α exposure, which may induce qualitatively distinct cellular responses at supraphysiological concentrations concentrations[1–4]. IFN-α2 at 1000 IU/mL was subsequently used for further experiments. In parallel, based on our dose-response experiments, 100 pM IFN-β was selected as the lowest concentration inducing maximal CD86 expression, with no additional response observed at higher concentrations. This choice was supported by previous studies showing potent biological effects of moderate IFN-β concentrations on human MoDCs, while higher concentrations may alter or inhibit specific DC functions [5-7]. Values are mean +/- SD of 9 independent measurements, except for the two higher doses of IFN-β (3 samples). Different symbols represent data from different HD samples.
